# Genome-wide characterization of MLO family in bread wheat shed light on its role in resistance to powdery mildew, biotic and abiotic stresses

**DOI:** 10.1101/2023.09.22.559087

**Authors:** Babar Hussain, Hamza Ramzan, Qasim Raza, Rana Muhammad Atif, Hikmet Budak, Zulfiqar Ali

## Abstract

Powdery mildew (PM) is a notorious disease that causes up to 62% of yield losses in wheat. The 66 PM resistance quantitative trait loci (QTL)/genes (Pm1 – Pm66) break down when new pathogen races interact with plants. The knockout of three wheat Mildew resistance locus o (MLO) has conferred stable resistance against PM. However, only seven MLO genes are known in bread wheat, which has limited the development of PM-resistant cultivars. Taking advantage of IWGSC Ref-seq v2.1, we identified 47 MLO genes in wheat that were distributed on all 21 chromosomes in a non-random fashion. Phylogenetic analysis showed that MLOs are divided into four clades/subfamilies, while clades I, II, III, and IV harbored 6, 28, 6, and 7 MLO genes, respectively. The phylogenetic patterns were strongly supported by gene structure and motif distribution in different clades. Motif analysis found 16 conserved motifs in wheat MLOs. Comparative phylogenetic tree of wheat, Arabidopsis, and rice MLOs classified the genes into four clades. Evolution analysis showed that segmental duplications and purifying selection are prevalent in TaMLOs. Finally, nine MLOs showed *in silico* expression in different tissues during growth and development. Eight genes (*TaMLO3/6-A2, TaMLO7-A1, TaMLO9-D1, TaMLO10-A2, TaMLO10-B1, TaMLO10-B2, TaMLO10-D1,* and *TaMLO10-D2*) showed overlapping expression under cold, drought, heat stress, and phosphate starvation. Several MLOs showed differential *in silico* expression under PM (*TaMLO10-A2, TaMLO3/6-A2, TaMLO7-A1*, *TaMLO10-D5*), stripe rust (*TaMLO9-D1*), and head blight (*TaMLO10-A2, TaMLO4-A1, TaMLO10-B1, TaMLO10-B2, TaMLO10-D1*, *TaMLO10-D2)*. The quantitative real-time polymerase chain reaction (qRT-PCR) showed that nine genes (*TaMLO3/6-A2*, *TaMLO3/6-B2*, *TaMLO3/6-D2*, *TaMLO8-A1*, *TaMLO8-B1*, *TaMLO8-D1*, *TaMLO9-A1*, *TaMLO10-A1*, and *TaMLO10-D1*) exhibited significant upregulation in PM-resistant line after 24, 48, and 72 hours of post-inoculation with the pathogen as compared with the susceptible cultivar. Whereas *TaMLO9-D1* showed downregulation. Thus, these MLO genes have a potential role in wheat under these conditions. Therefore, we hope that the MLO genes identified in this study will be edited through CRISPR/Cas9 and/or will be overexpressed to develop PM and disease-resistant and abiotic stress-tolerant wheat.

## 1. Introduction

Climate change not only alters temperature and rain patterns, but it also results in the emergence of new pathogen races. Several fungal diseases such as leaf rust, stripe rust, stem rust, smuts, septoria leaf blotch, fusarium head blight (FHB), and powdery mildew (PM) (Hussain et al. 2022; Öztürk et al. 2025). PM caused by the fungus *Blumeria graminis* is a devastating disease that is prevalent globally and causes up to 62% of yield losses in wheat (Singh et al. 2016). Therefore, efforts have been made to identify resistance genes for PM. A total 66 resistance genes (designated as *Pm1*-*Pm66*) have been mapped in wheat after decades-long efforts (McIntosh et al. 2019). Ten of these genes (*Pm1a, Pm2, Pm3, Pm3b, Pm5e, Pm21, Pm24, Pm38/Lr34/Yr18/Sr57*, and *Pm41, Pm46/Lr67/Yr46/Sr55, Pm60*) have been cloned in wheat to develop PM resistance (Hussain et al. 2022). It should be noted that these are not MLO genes but are diverse resistance genes identified through quantitative trait loci (QTL) mapping. However, cloning techniques such as map-based cloning are difficult to perform, tedious, and time consuming and need specialized lab facilities and skilled manpower. Therefore, the process to develop PM resistance is slow against fast-evolving pathogens. Additionally, resistance against pathogens breaks down when new pathogen races interact with host plants (Hussain 2015).

In barley, loss of function mutations in a Mildew resistance locus o (Mlo) gene conferred a broad spectrum resistance against PM. This resistance was so durable that it lasted in the field for more than 30 years (Jørgensen 1992). The loss-of-function mutations yielded pleiotropic effects such as necrotic leaf spotting and reduced grain yield, but those were overcome by breeding efforts. The stable nature of MLO-based resistance has resulted in a great interest in these genes. Till date, genome-wide identification of MLO genes has been performed in many plant species including *Arabidopsis thaliana* (Devoto et al. 2003), *N. benthamiana* (Kusch et al. 2016), *B. distachyon* (Ablazov and Tombuloglu 2016), rice (Liu and Zhu 2008; Nguyen et al. 2016), foxtail millets and barley (Kusch et al. 2016), maize (Gong et al. 2025), oat (Reilly et al. 2021), tomato (Zheng et al. 2016; Shi et al. 2020), apple and peach (Kusch et al. 2016), grapevine (Zheng et al. 2016), diploid and octaploid strawberry (Tapia et al. 2021), several Rosaceae species (Tian et al. 2022), lentil (Polanco et al. 2018), chickpea, mung bean, pea, groundnut, narrow-leaf lupin, pigeon pea, soybean and common bean (Rispail and Rubiales 2016), bitter gourd (Chen et al. 2021), bottle gourd, wax gourd, watermelon, melon, cucumber, kaffir lime, pumpkin, and crookneck pumpkin (Gong et al. 2025) among many other species.

MLO genes are well known for susceptibility to the PM disease in plants, and their loss-of-function mutations have been generated to create PM-resistant plants. However, only seven MLO genes were known in bread wheat to date (Konishi et al. 2010), largely due to the non-availability of its reference genome. The *TaMLO-A1*, *TaMLO-B1*, and *TaMLO-D1* genes were knocked out simultaneously in bread wheat using transcription activator–like effector nuclease (TALEN), and clustered regularly interspaced short palindromic repeats-CRISPR/Cas9 system, which conferred heritable broad-spectrum resistance against PM (Wang et al. 2014). Similarly, simultaneous knockout of these genes through Targeting Induced Local Lesions IN Genomes (TILLING) enhanced PM resistance without pleiotropic phenotypes (Acevedo Garcia et al. 2017). Recently, CRISPR/Cas9 was used to introduce a 304-kilobase pair deletion in *TaMLO-B1* that conferred robust PM resistance without pleiotropic effects. This was due to altered chromatin that led to the ectopic activation of *tonoplast monosaccharide transporter 3* (*TaTMT3B*) and negated the negative effects of MLO disruption (Li et al. 2022). Additionally, simultaneous mutations in above-mentioned genes provided highly effective PM resistance without pleiotropic effects (Ingvardsen et al. 2023).

The newly emerged CRISPR/Cas system has been used to edit the effector-binding elements of susceptibility genes (Hussain et al. 2018; Hussain and Ahmad 2022) to confer durable and broad-spectrum resistance against diseases. Keeping in view the availability of limited MLOs in the bread wheat, we performed a genome-wide search to identify all MLO genes in bread wheat and studied their evolution through phylogenetic and duplication analysis. Furthermore, gene structure and motif analysis were performed to study the diversity in MLO genes. We also performed MLO expression under different growth and development stages, PM, and other biotic and abiotic stresses through an *in-silico* approach and under PM by qRT-PCR. We hope that additional MLO genes identified in this study, especially the ones showing differential expression under different conditions, will be edited/knocked out through CRISPR/Cas9 and other mutation-inducing techniques to improve PM, biotic, and abiotic stress tolerance in wheat.

## 2. Materials and Methods

### 2.1. Sequence Retrieval and Genome-Wide Identification of MLO Genes

The *T. aestivum* RefSeq v2.1 genome assembly, proteome, and transcriptome, along with the general feature format file (GFF3), were downloaded from the website (https://urgi.versailles.inrae.fr/download/iwgsc/IWGSC_RefSeq_Assemblies/v2.1/). A total of 15 *A. thaliana* MLO sequences were downloaded from The Arabidopsis Information Resource (TAIR) database (https://www.arabidopsis.org/ ). The MLO sequences of rice MLOs were obtained from a previous study (Liu and Zhu 2008). Genome-wide identification of MLO genes in the *Triticum aestivum* genome was performed through the Basic Local Alignment Search Tool (BLAST) function of TB tools (Chen et al. 2020). For the purpose, *A. thaliana* MLO sequences were used as query sequences. Additionally, an HMMER-based search was performed to predict the MLO proteins in wheat genomes for further confirmation (Potter et al. 2018). The presence of a complete MLO domain in wheat proteins was confirmed through the NCBI Conserved Domain Database (CDD) search tool v3.19 (https://www.ncbi.nlm.nih.gov/Structure/cdd/wrpsb.cgi) (Wang et al. 2023) and Pfam/Interpro (http://pfam.xfam.org/). The proteins without a complete MLO conserved domain were not considered for further analysis, as such genes lose their functions (Raza et al. 2022).

Moreover, Expasy server’s ProtParam tool (https://web.expasy.org/protparam/) was used for the estimation of molecular weight and isoelectric point of the MLO proteins. The subcellular localization of MLO proteins was predicted with a web-based server, Bologna Unified Subcellular Component Annotator (BUSCA) (https://busca.biocomp.unibo.it/). For the purpose, the organ that proclaimed the highest value was taken as the location of the protein.

### 2.2. Chromosomal Mapping of MLO Genes

The identified genes were mapped on the chromosomes by using the graphics interface of TB Tools software (Chen et al. 2020). The chromosomal locations of the MLO genes were extracted from the GFF3 file.

### 2.3. Phylogenetic Analysis

The wheat MLO sequences were aligned through multiple sequence alignment (MSA) tool, i.e., Clustal Omega (https://www.ebi.ac.uk/jdispatcher/msa/clustalo) (Sievers et al. 2011), with default settings. The resultant multiple sequence alignment was used to construct a phylogenetic tree with the maximum likelihood (ML) method using MEGA v11.0 (Tamura et al. 2021). For the purpose, the Jones-Taylor-Thornton (JTT) model for amino acid substitution was utilized. This model is the standard for constructing maximum likelihood trees from amino acid/protein sequences. For statistical replication, 1,000 replications for the bootstrapping were used, i.e., the software assessed the reliability of each node/branch by resampling the protein data and reconstructed the phylogenetic tree 1,000 times. After the completion of 1,000 bootstraps, the tree was exported in the form of Newick, and the tree was edited for better visualization with the iTol program (https://itol.embl.de/).

For comparative phylogenetic analysis, 47 TaMLOs identified in this study, along with 15 *AtMLOs* (Devoto et al. 2003) and 12 rice MLO (Liu and Zhu 2008; Nguyen et al. 2016) protein sequences, were aligned through Clustal Omega. The resultant MSA was used to construct the tree through the IQ-TREE program (Trifinopoulos et al. 2016).

### 2.4. Gene Structure and Motif Analysis

The genomic and coding sequences (CDS) of all *TaMLO* genes were pasted in the Gene Structure Display Server (GSDS) tool (http://gsds.gao-lab.org/) in FASTA format to visualize the exons, introns, and down/upstream regions of the genes. The conserved motif analysis was performed by the MEME 5.1 tool (http://meme-suite.org/tools/meme). We tried different settings for the proper identification of conserved motifs and found a suitable setting of 16 as the maximum number of motifs and 50 as the maximum length of motifs. The motif data was downloaded as an XML file, and TB tools software (Chen et al. 2020) was used to visualize the gene structure. All motif sequences were searched in the Simple Modular Architecture Research Tool (SMART) database (http://smart.embl-heidelberg.de/) to find the known domains/repeats/motifs. Afterwards, complete MLO protein sequences were searched in Deep Transmembrane Helices Hidden Markov Models (TMHMM)-1.0 server (https://services.healthtech.dtu.dk/services/DeepTMHMM-1.0/) to predict the number of trans-membranes in MLO proteins. The complete protein sequences were pasted in the Expasy Protparam server (https://web.expasy.org/protparam/) to calculate the leucine percentage in all MLO genes.

### 2.5. Divergence (Gene Duplication) and MicroRNA Analysis

We followed a previously described procedure (Raza et al. 2022) for the purpose. The CDS of all *TaMLO* genes were aligned using Clustal Omega (https://www.ebi.ac.uk/jdispatcher/msa/clustalo) (Sievers et al. 2011). The resulting MSA was MAFFT aligned (pairwise) in all possible combinations using Sequence Demarcation Tool v1.2 (Muhire et al. 2014) to calculate the sequence identities. Gene pairs with at least 90% identity (E value < 1e^−10^) were classified as duplicates. The duplicated gene pairs that were mapped on the same chromosome were considered as tandem duplication. The gene pairs located on different chromosomes were defined as segmental duplication. The CDS of duplicate genes were MAFFT aligned and subjected to Synonymous Non-Synonymous Analysis Program v2.1.1 (www.hiv.lanl.gov/content/sequence/SNAP/SNAP.html) to compute the synonymous (Ks) and non-synonymous (Ka) substitutions. The Ka/Ks was calculated to predict the codon selection during the evolution. An approximate divergence time for the duplicated genes was calculated by using the formula T = Ks/2r × 10−6, assuming a substitution rate (r) of 6.5 × 10−9 substitutions/synonymous site/year (El Baidouri et al. 2017).

For microRNA (miRNAs) identification, the transcript sequences of identified TaMLOs were searched on the psRNATarget server (https://www.zhaolab.org/psRNATarget/) to predict micro-RNAs potentially targeting the identified genes.

### 2.6. *In silico* Expression Analysis

The RNA-seq expression data for TaMLO genes during growth and development under different abiotic (cold, drought, heat, and phosphate starvation) and biotic (FHB, PM, and stripe rust) stresses were retrieved from the expVIP Wheat Expression Browser as Log2 Transcripts Per Million (TPM) (Borrill et al. 2016; Ramírez-González et al. 2018). The expression heatmaps were generated with TB tools (Chen et al. 2020).

### 2.7. Quantitative PCR Analysis

#### 2.7.1. Plant Materials, Growth Conditions, and Stress Treatments

A PM-resistant wheat line, N9134, that shows high resistance to all *Blumeria graminis* f. sp. tritici (Bgt) races in China (Zhang et al. 2014), along with a susceptible wheat cultivar, Fielder, were selected for quantitative real-time polymerase chain reaction (qRT-PCR)-based expression analysis. Seeds of both genotypes were obtained from Peking University Institute of Advanced Agricultural Sciences, China. Healthy seeds of both genotypes were disinfected (0.1% HgCl_2_ for 10 minutes), rinsed with autoclaved distilled water, surface dried on filter paper, sown in an autoclaved soil + peatmoss mixture, and kept under controlled conditions (22 ± 1 °C, 60 ± 5% relative humidity,16 h/8 h light and dark cycle). Two-weeks-old seedlings with consistent growth were inoculated with Bgt conidia. The control seedlings were grown without Bgt inoculation. The control and inoculated leaves were harvested at 0, 24, 48, and 72 hours post inoculation (hpi). Leaf samples from 3 independent seedlings were collected, flash frozen in liquid nitrogen and kept at - 80 °C until RNA extraction.

#### 2.7.2. RNA Extraction and qRT-PCR Analysis

The total RNA was extracted using the SPARKeasy RNA extraction kit (SparkJade, China) by following manufacturer instructions. After confirming RNA quality through agarose gel electrophoresis, reverse transcription was performed using the SPARKscript II One Step RT-PCR kit. Gene-specific primers were designed through the NCBI Primer-BLAST tool, and qRT-PCR was performed using ChamQ Blue Uninversal SYBR qPCR Master Mix (Vazyme, China) on a QuantStudio^TM^ 5 system (Thermo Fisher Scientific). The 20 µL reaction system included 10 µL SYBR mix, 0.4 µL of each primer pair, 5 µL diluted cDNA, and 4.2 µL RNase-free water. For internal control, TaActin (TraesCS5D02G132200) was used. The qPCR test procedure was as follows: 95 °C for 1 minute, followed by 40 cycles of 95 °C for 10 s and 60 °C for 30 s. The results were analyzed using the 2−ΔΔCt method (Livak and Schmittgen 2001). The analysis was carried out with three biological repeats, and mean values were used for generating graphs. A list of primer pairs for MLO genes and internal control is provided in **Table S1**.

## 3. Results and Discussion

### 3.1. Genome-Wide Identification and Properties of MLO Genes

The bread wheat genome was sequenced after 13 years of work due to its large size (> 17 Gb) (Hussain et al. 2022). This has greatly limited the discovery and characterization of MLOs in wheat, and only seven of them (Konishi et al. 2010) were reported. Utilizing the bread wheat RefSeq v2.1, we initially identified 61 MLOs in wheat. For the purpose, *A. thaliana* MLO proteins were used as query sequences in the BLAST function of TB tools. Among these, two duplicate genes were deleted by sorting the genes by their names. The MSA showed that four genes contained incomplete MLO domains (pseudogenes). As pseudogenes lose their function (Panchy et al. 2016), so they were deleted from further analysis. Eight genes were found to be isomers in the phylogenetic tree, so they were also deleted; thus, we got 47 non-redundant MLO genes **(Table 1)**. The presence of the MLO domain in these proteins was validated through NCBI CDD search and the Pfam database.

**Table 1.** Gene IDs, newly assigned names and properties of identified MLO genes in *Triticum aestivum* L.

| S# | Gene IDs | New Gene Names | Chr. | Subfamily | Gene Start | Gene End | Gene Size (bp) | Strand |
| --- | --- | --- | --- | --- | --- | --- | --- | --- |
| 1 | TraesCS1A03G0539600 | *TaMLO1-A1* | 1A | III | 364505363 | 364509878 | 4515 | - |
| 2 | TraesCS1B03G0628500 | *TaMLO1-B1* | 1B | III | 398073082 | 398077619 | 4537 | - |
| 3 | TraesCS1D03G0519600 | *TaMLO1-D1* | 1D | III | 293427154 | 293431832 | 4678 | - |
| 4 | TraesCS3A03G0944700 | *TaMLO2-A1* | 3A | II | 648479170 | 648482130 | 2960 | - |
| 5 | TraesCS3B03G1076900 | *TaMLO2-B1* | 3B | II | 689913560 | 689916559 | 2999 | + |
| 6 | TraesCS3D03G0877800 | *TaMLO2-D1* | 3D | II | 513351507 | 513355165 | 3658 | - |
| 7 | TraesCS5A03G1158600 | *TaMLO3/6-A1* | 5A | IV | 664743006 | 664756436 | 13430 | + |
| 8 | TraesCS5A03G1159400 | *TaMLO3/6-A2* | 5A | IV | 664797864 | 664801136 | 3272 | - |
| 9 | TraesCS5A03G1150400 | *TaMLO3/6-A3* | 5A | IV | 661126976 | 661129967 | 2991 | - |
| 10 | TraesCS4B03G0835400 | *TaMLO3/6-B1* | 4B | IV | 611711110 | 611728680 | 17570 | + |
| 11 | TraesCS4B03G0836900 | *TaMLO3/6-B2* | 4B | IV | 612012622 | 612014655 | 2033 | - |
| 12 | TraesCS4D03G0743900 | *TaMLO3/6-D1* | 4D | IV | 483173859 | 483176970 | 3111 | + |
| 13 | TraesCS4D03G0744500 | *TaMLO3/6-D2* | 4D | IV | 483205801 | 483208560 | 2759 | - |
| 14 | TraesCS2A03G1299000 | *TaMLO4-A1* | 2A | **I** | 768633437 | 768638338 | 4901 | - |
| 15 | TraesCS2B03G1489500 | *TaMLO4-B1* | 2B | **I** | 788988904 | 788994031 | 5127 | - |
| 16 | TraesCS2D03G1262400 | *TaMLO4-D1* | 2D | **I** | 638659281 | 638665452 | 6171 | - |
| 17 | TraesCS2A03G0775400 | *TaMLO5-A1* | 2A | **II** | 539693546 | 539696928 | 3382 | - |
| 18 | TraesCS2B03G0856800 | *TaMLO5-B1* | 2B | **II** | 479483631 | 479486919 | 3288 | + |
| 19 | TraesCS2D03G0715400 | *TaMLO5-D1* | 2D | **II** | 400923755 | 400926943 | 3188 | - |
| 20 | TraesCS7A03G0928000 | *TaMLO7-A1* | 7A | **II** | 561794230 | 561801544 | 7314 | - |
| 21 | TraesCS7B03G0775700 | *TaMLO7-B1* | 7B | **II** | 524484720 | 524491473 | 6753 | - |
| 22 | TraesCS7D03G0891400 | *TaMLO7-D1* | 7D | **II** | 493179947 | 493187259 | 7312 | - |
| 23 | TraesCS6A03G0387200 | *TaMLO8-A1* | 6A | **II** | 154203217 | 154207538 | 4321 | + |
| 24 | TraesCS6B03G0494800 | *TaMLO8-B1* | 6B | **II** | 232292016 | 232296365 | 4349 | - |
| 25 | TraesCS6D03G0340700 | *TaMLO8-D1* | 6D | **II** | 148637851 | 148642268 | 4417 | - |
| 26 | TraesCS1A03G0668200 | *TaMLO9-A1* | 1A | **II** | 457568166 | 457572624 | 4458 | - |
| 27 | TraesCS3A03G0013800 | *TaMLO9-A2* | 3A | **II** | 6965742 | 6973314 | 7572 | - |
| 28 | TraesCS1B03G0759600 | *TaMLO9-B1* | 1B | **II** | 484447774 | 484452526 | 4752 | - |
| 29 | TraesCS1D03G0630400 | *TaMLO9-D1* | 1D | **II** | 357744682 | 357749428 | 4746 | - |
| 30 | TraesCS3D03G0010400 | *TaMLO9-D2* | 3D | **II** | 1805922 | 1810130 | 4208 | - |
| 31 | TraesCS2A03G0163000 | *TaMLO10-A1* | 2A | **II** | 42897580 | 42900950 | 3370 | - |
| 32 | TraesCS2A03G0163100 | *TaMLO10-A2* | 2A | **II** | 42905631 | 42908828 | 3197 | - |
| 33 | TraesCS6A03G0805500 | *TaMLO10-A3* | 6A | **II** | 545643614 | 545648358 | 4744 | - |
| 34 | TraesCS2B03G0227100 | *TaMLO10-B1* | 2B | **II** | 65216529 | 65219835 | 3306 | - |
| 35 | TraesCS2B03G0227200 | *TaMLO10-B2* | 2B | **II** | 65315357 | 65318657 | 3300 | - |
| 36 | TraesCS6B03G0955000 | *TaMLO10-B3* | 6B | **II** | 601764897 | 601769827 | 4930 | - |
| 37 | TraesCS2D03G0160400 | *TaMLO10-D1* | 2D | **II** | 37249636 | 37253058 | 3422 | - |
| 38 | TraesCS2D03G0160500 | *TaMLO10-D2* | 2D | **II** | 37272260 | 37275524 | 3264 | - |
| 39 | TraesCS6D03G0681600 | *TaMLO10-D3* | 6D | **II** | 418198229 | 418205340 | 7111 | - |
| 40 | TraesCS7D03G0041500 | *TaMLO10-D4* | 7D | **II** | 8641641 | 8683479 | 41838 | - |
| 41 | TraesCS7D03G0062000 | *TaMLO10-D5* | 7D | **II** | 14007248 | 14014362 | 7114 | + |
| 42 | TraesCS4A03G0580300 | *TaMLO11-A1* | 4A | **I** | 518674297 | 518679167 | 4870 | + |
| 43 | TraesCS4B03G0222000 | *TaMLO11-B1* | 4B | **I** | 104220694 | 104226387 | 5693 | + |
| 44 | TraesCS4D03G0182500 | *TaMLO11-D1* | 4D | **I** | 69109133 | 69114152 | 5019 | + |
| 45 | TraesCS5A03G0963900 | *TaMLO12-A1* | 5A | **III** | 599592378 | 599596257 | 3879 | + |
| 46 | TraesCS5B03G1014800 | *TaMLO12-B1* | 5B | **III** | 590155469 | 590160172 | 4703 | + |
| 47 | TraesCS5D03G0918700 | *TaMLO12-D1* | 5D | III | 481711884 | 481717428 | 5544 | + |

Following the confirmation, renaming of wheat MLOs was done according to their localization with rice orthologs in the comparative phylogenetic tree (**Figure 3**). For example, three MLO genes that occurred in close proximity to *OsMLO1* were renamed as *TaMLO1-A1*, *TaMLO1-B1*, and *TaMLO1-D1* following the standard guidelines for wheat gene families (Boden et al. 2023). Here, A, B, and D represent the orthologs of *OsMLO1* present in the A, B and D genomes of wheat, while the 1 following these letters shows the first wheat MLO in the respective sub-genome (**Table 1**).

A comparison of two previous studies shows that there is no relationship between genome size and the number of MLO genes in a species (Kusch et al. 2016; Shi et al. 2020). For example, maize (genome size of 2,400 Mb) has a lesser number of MLO proteins (13) than the 15 MLOs in *A. thaliana* with a 120 Mb genome. On the other hand, a gymnosperm, Norway spruce (*Picea abies*), with a genome size of 20 Gb, has only 14 MLO proteins. This is thought to be linked with large-scale occurrences of gene losses, gene expansion, and indels during plant evolution (Shi et al. 2020).

However, based on the previous studies and our analysis, we infer that the number of MLO genes in a species does depend upon its ploidy level and not the genome size. Many diploid plant species have 11 to 17 MLOs. For example, 11 MLOs in *B. distachyon*, oat, and barley; 12 MLOs in rice and foxtail millet; 13 ZmMLOs in maize; 14 CsMLOs in cucumber; 15 MLOs in *A. thaliana* and lentil; 16 SlMLOs in tomato; and 17 VvMLOs in grapevine (Devoto et al. 2003; Kusch et al. 2016; Nguyen et al. 2016; Polanco et al. 2018; Shi et al. 2020; Reilly et al. 2021) were reported. This number doubles for tetraploid species; e.g., *N. benthamiana* has 26 (Kusch et al. 2016), and soybean has 31 MLOs (Rispail and Rubiales 2016). Our study showed that hexaploid bread wheat has 47 and octaploid strawberry has 68 MLOs (Tapia et al. 2021). Therefore, it can be concluded that the number of MLOs in tetraploid, hexaploid, and octaploid plants is two, three, and four times greater than in the diploid plants.

The isoelectric point values varied from 7.36 to 9.87, while the molecular weight of the proteins was in the range of 46.98 to 68.50 kDa. The peptide length of these MLO proteins ranged from 409 to 612 amino acids, and the average length was 514 amino acids. For any wheat gene, its three orthologs present in A, B, and D genomes have almost the same number of amino acids and protein molecular weight (**Table S2**). This range of 400 to 600 amino acids in MLO proteins seems quite conserved in plants, as it has been reported in 28 plant species in a previous study (Shi et al. 2020). The subcellular localization of MLO proteins through the BUSCA tool revealed that all MLOs were located on the cell membrane (**Table S2**). This has been reported previously (Kusch et al. 2016; Shi et al. 2020); thus, their function may be linked to membrane signal transduction and cell-to-cell communication.

### 3.2. Distribution of MLOs in Bread Wheat Genome/s

None of the seven wheat MLO genes that were identified in a previous study (Konishi et al. 2010) were mapped on any chromosome. This was due to the non-availability of a reliable wheat reference genome till 2018 (Appels et al. 2018). The reference genome has enabled us to scan the whole genome, identify all 47 wheat MLO genes, and assign them to all 21 chromosomes for the first time. A total of 16, 14, and 17 MLO genes were mapped on the A, B, and D genomes of wheat, respectively, in a non-random fashion (Table 1; **Figure 1A, 1B, 1C**). Such a non-random distribution of plant gene families originates from long terminal repeat retrotransposons through evolution (Gao et al. 2004) and has been reported in other plant species as well (Ahmed et al. 2021). A previous study showed that 174 MLO genes in 18 species were also distributed on specific chromosomes or in a non-random pattern (Shi et al. 2020).

**Figure 1A:**
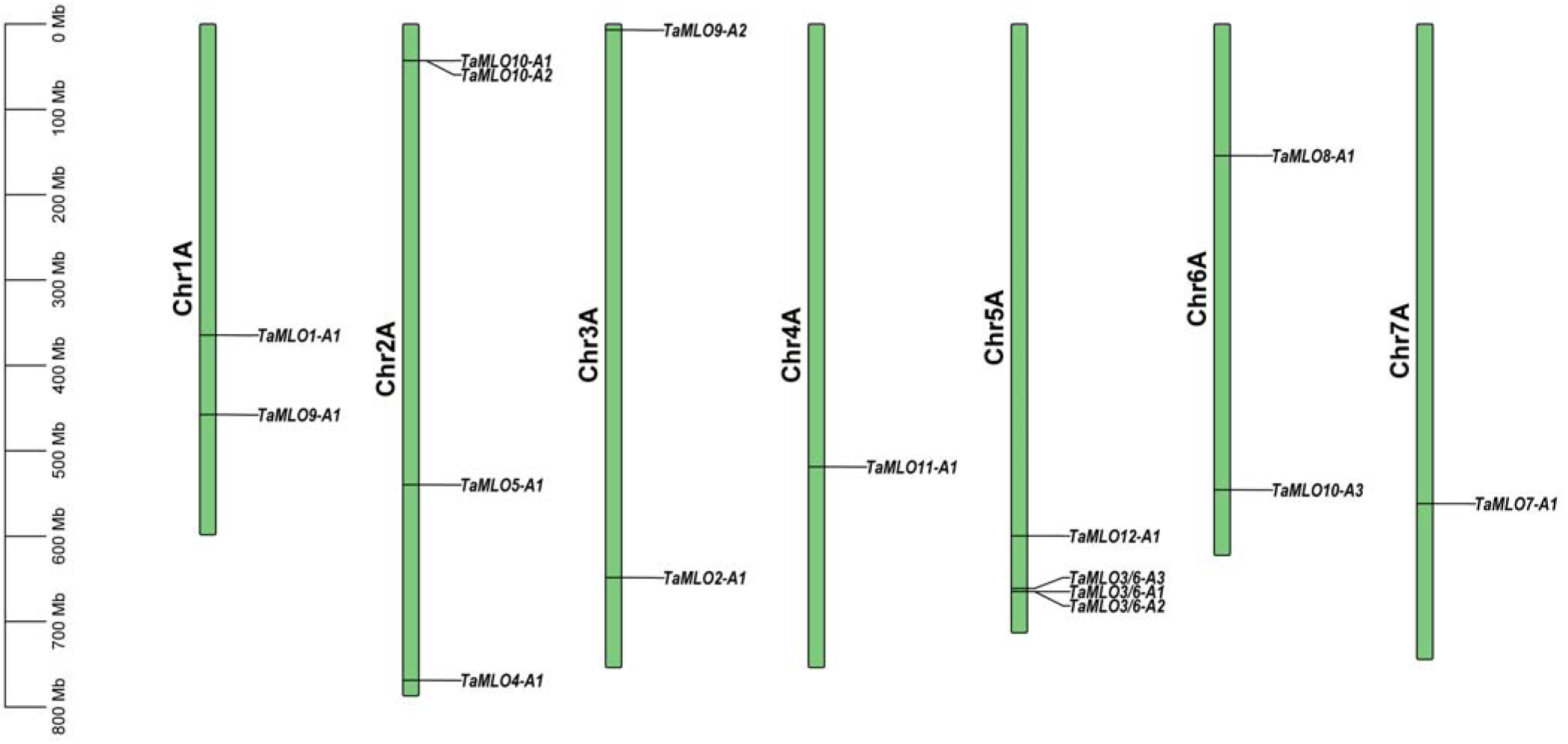
Chromosomal mapping of 16 TaMLO genes on AA genome of bread wheat

**Figure 1B:**
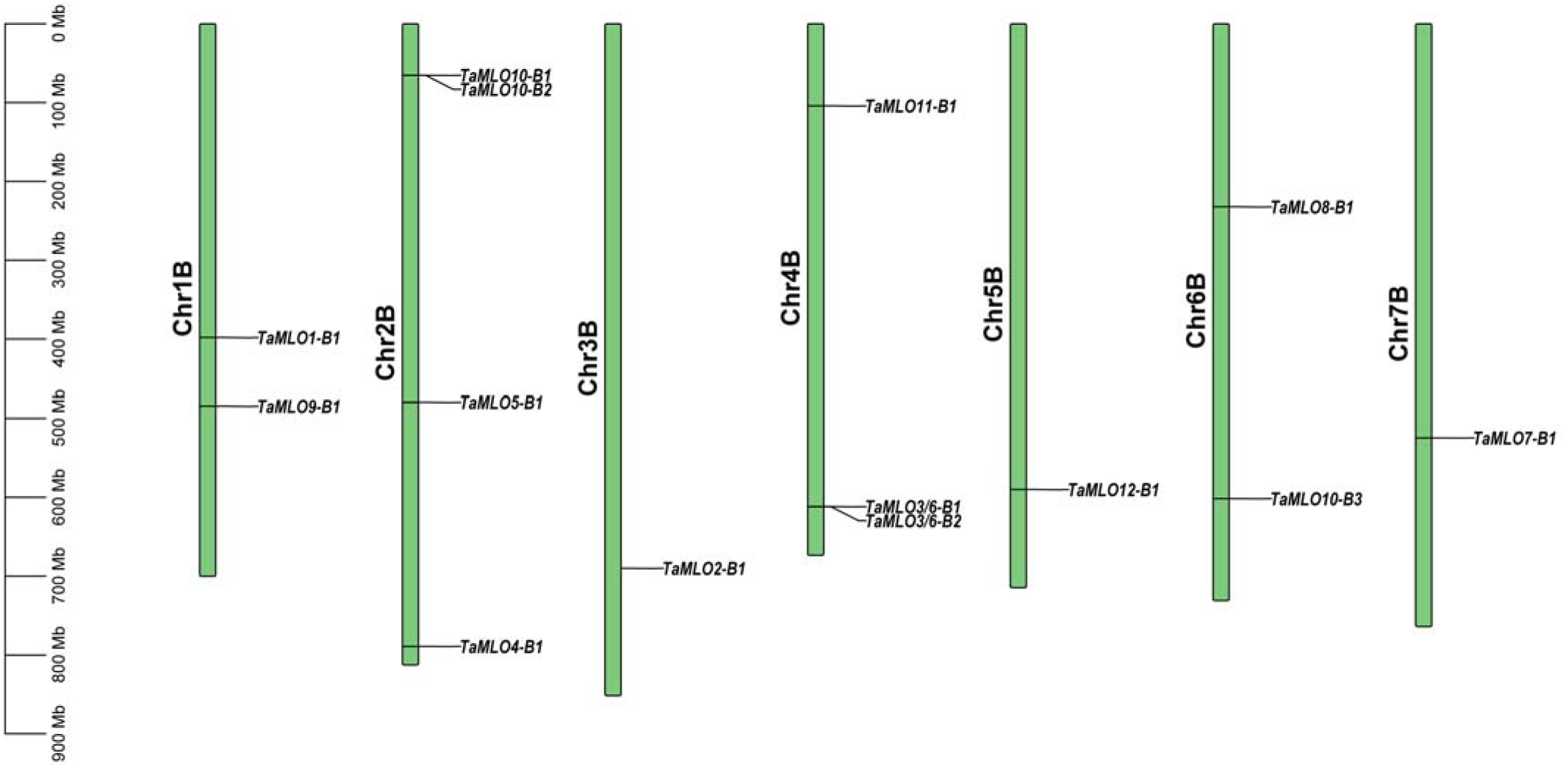
Chromosomal mapping of 14 TaMLO genes on BB genome of bread wheat

**Figure 1C:**
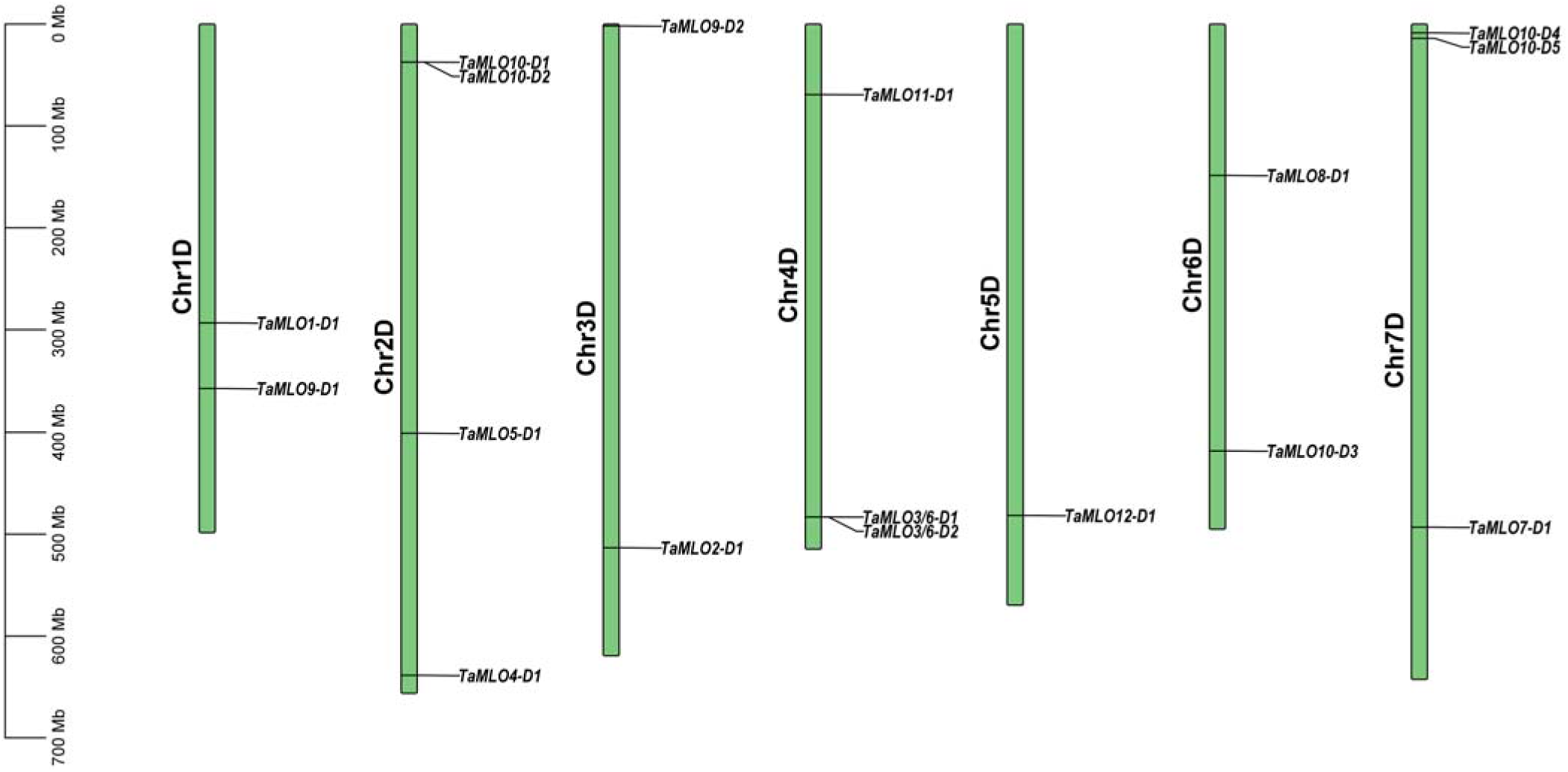
Chromosomal mapping of 17 TaMLO genes on DD genome of bread wheat

The chromosomes 2A, 5A, 2B, and 2D harbored the maximum four TaMLO genes each, while chromosomes 1A, 1B, 1D, 6A, 6B, and 6D contained two genes each. Interestingly, chromosome 5A harbored four TaMLO genes, while 5B and 5D contained only one gene. Similarly, chromosome 4A harbored only one gene in contrast to three TaMLO genes on both 4B and 4D. Additionally, the number of mapped genes on 7D was three in comparison to one gene on 7A and 7B, while two, one, and two genes were mapped on chromosomes 3A, 3B, and 3D, respectively (**Figure 1A, 1B, and 1C**). Such varying numbers of MLO genes on different chromosomes of a species were also reported in rice, soybean, and tomato (Liu and Zhu 2008; Shi et al. 2020). Additionally, we observed clustering of MLO genes on chromosomes 2A, 5A, 2B, 4B, 2D, and 4D (**Figure 1A, 1B, 1C**). Such clustering was not observed in diploid rice (Liu and Zhu 2008) but was observed in tetraploid soybean (Rispail and Rubiales 2016) and octaploid strawberry (Tapia et al. 2021). In conclusion, the distribution of MLO genes on chromosomes is very specific and follows a non-random and conserved pattern across different species.

### 3.3. Phylogenetic Analysis of MLO Genes

We constructed a maximum likelihood tree to demonstrate the evolutionary relationship among the MLO proteins in modern wheat. The phylogenetic tree showed the distribution of MLO proteins into four distinct subfamilies or clades (**Fig. 2a**). The subfamily or clade numbers were assigned by following the clade numbering in *A. thaliana* (Devoto et al. 2003), as per standard practice. The distribution of 47 MLO superfamily genes into four subfamilies is because of group-specific motifs that are conserved among the proteins (**Fig. 2a, c**). Subfamily II represents the largest clade with 28 members (59.57% TaMLOs), while subfamilies I, III, and IV have six, six, and seven members, respectively. Subfamily I contained *TaMLO4-A1, TaMLO4-B1, TaMLO4-D1, TaMLO11-A1, TaMLO11-B1, and TaMLO11-D1*. The subfamily III has *TaMLO1-A1*, *TaMLO1-B1*, *TaMLO1-D1*, *TaMLO12-A1*, *TaMLO12-B1*, and *TaMLO12-D1*, while subfamily IV has seven MLOs, i.e., *TaMLO3/6-A1, TaMLO3/6-A2, TaMLO3/6-A3, TaMLO3/6-B1, TaMLO3/6-B2, TaMLO3/6-D1, and TaMLO3/6-D2*. Additionally, 28 MLOs in clade II were further divided into three subclades, i.e., IIA, IIB, and IIC (**Fig. 2a**). More than 50% of the MLO proteins in many monocot species were classified into clade II (Kusch et al. 2016), while clade IV either has a very low number of MLOs or is completely lost in some dicot species.

**Figure 2:**
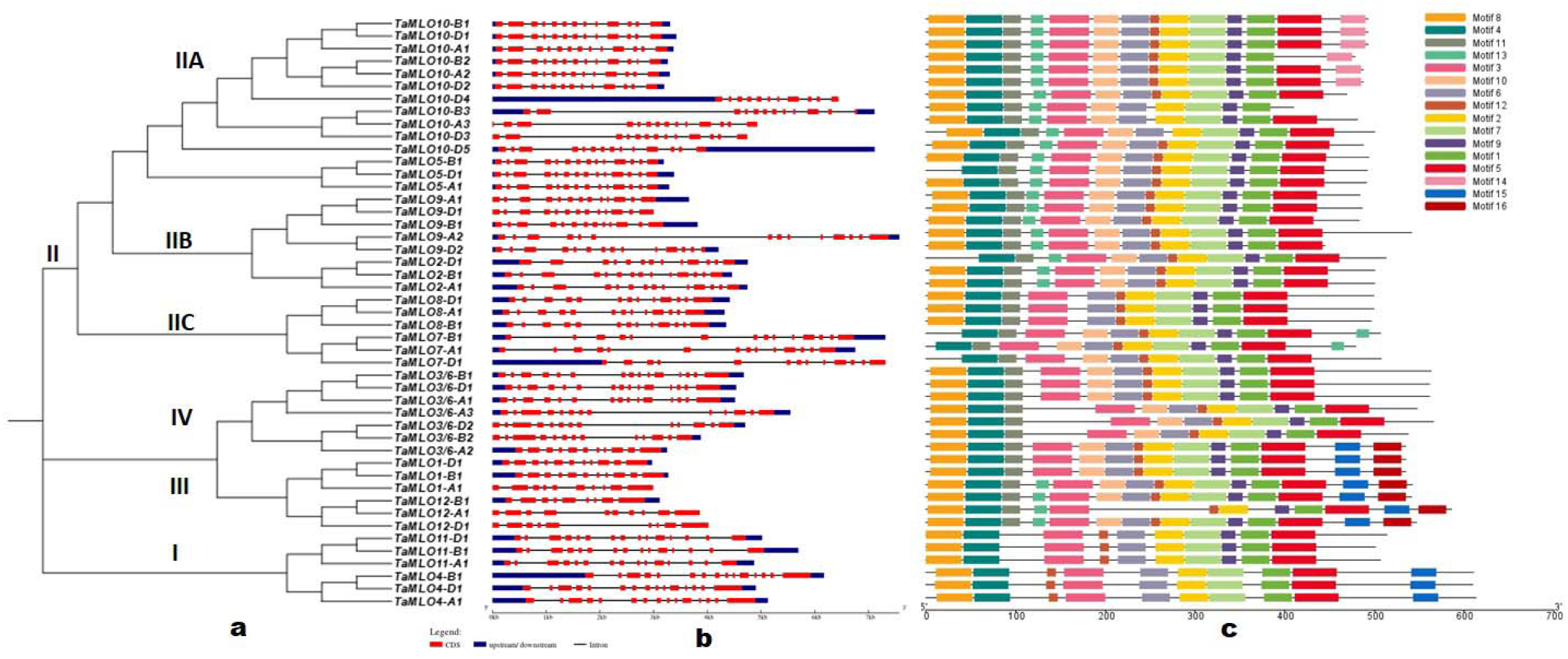
Comparison of a) the maximum likelihood phylogenetic tree of 47 TaMLO proteins constructed with 1000 bootstrap replications, with b) the gene structure of full-length genomic sequences of 47 MLOs, wred boxes represent the exons, and lines connecting the exons show the introns, while blue boxes represent the upstream and downstream sequences of the genes, and c) the 16 conserved motifs located on the 47 MLO proteins.

**Figure 3:**
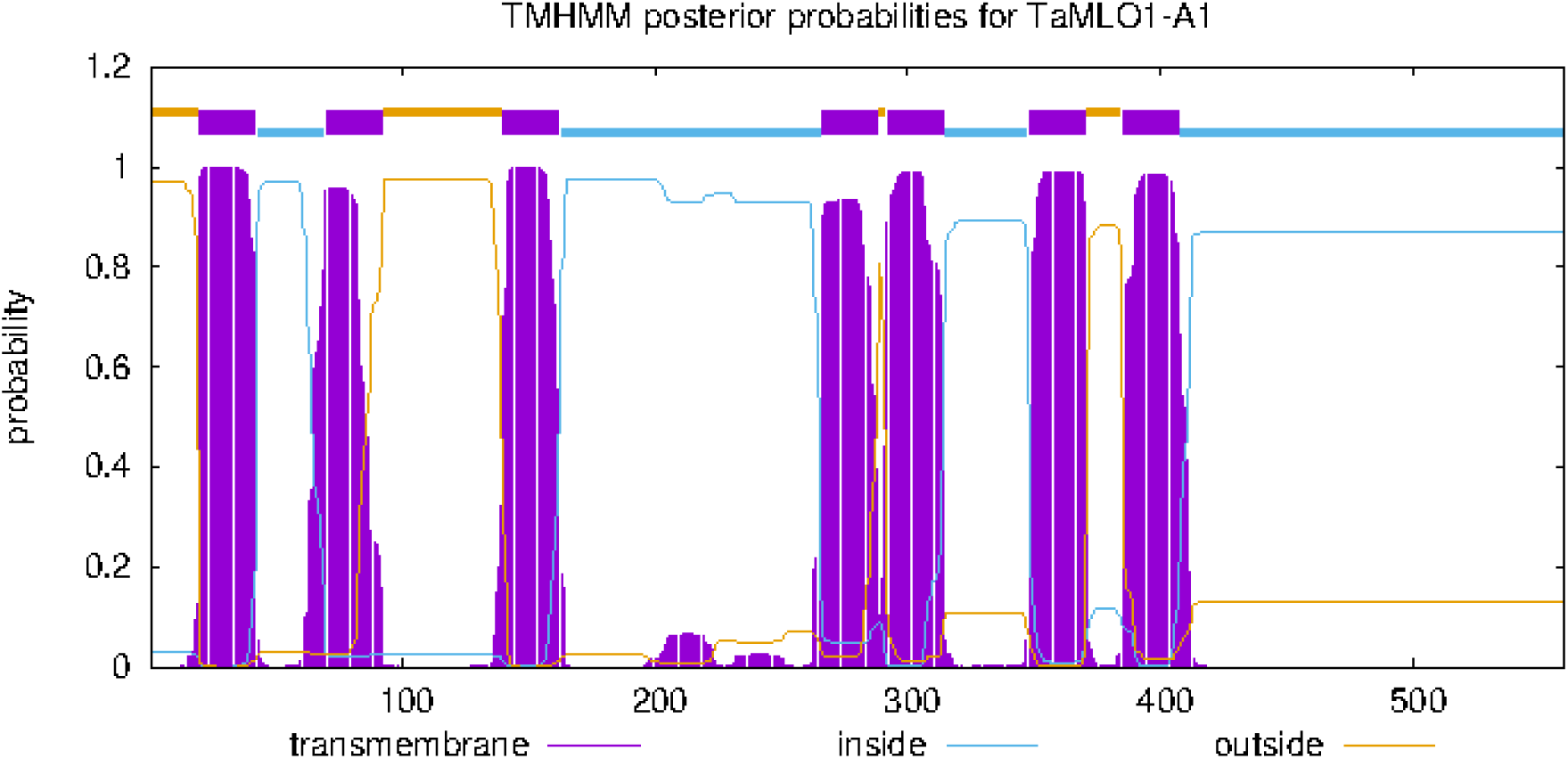
The graphical representative of TMHMM depicting the posterior probabilities for transmembrane, outside (exterior), and inside (cytoplasm) regions. In this example (TaMLO1-A1), server predicted six transmembrane-TM regions.

Seven TaMLO genes were also classified into four subfamilies (Konishi et al. 2010). Similarly, MLOs in other monocots such as *B. distachyon*, rice, maize, oat, barley, and foxtail millet were also divided into four clades (Liu and Zhu 2008; Kusch et al. 2016; Nguyen et al. 2016; Reilly et al. 2021). However, this highly conserved pattern of four MLO subfamilies is exclusive to monocots. A basal angiosperm, Amborella, has six clades for MLOs. It is believed to have diverged from the other angiosperms more than 160 million years ago (MYA), before the split of monocots and dicots (120 MYA). Therefore, it is possible that six clades for MLOs were also present in the ancestors of the monocots, but clades V and VI were lost during the monocot evolution (Kusch et al. 2016). Additionally, MLOs for several dicots (*A. thaliana*, soybean, cucumber, apple, wild strawberry, tomato, peach, grapevine, etc.) were also divided into six subfamilies (Devoto et al. 2003; Kusch et al. 2016). A comprehensive phylogenetic analysis concluded that MLO proteins in dicots should divide into seven clades (I–VII). However, most dicots lost their MLOs in clade IV or VII during the evolution (Kusch et al. 2016), thus explaining the conserved division of MLO proteins into six clades in dicots.

The subfamily distribution of MLOs is associated with their conserved functions. Many eudicot MLOs found in clade V are involved in PM susceptibility. For example, genes governing PM susceptibility in *A. thaliana (AtMLO2*, *AtMLO6*, and *AtMLO12)*; pea (*Er1*/*PsMLO1);* tomato (*SlMLO1*); grapevine (*VvMLO3* and *VvMLO4);* cucumber (*CsaMLO1*, *CsaMLO8*, and *CsaMLO11*); and lentil (*LcMLO1* and *LcMLO3*) are found in clade V (Bai et al. 2008; Humphry et al. 2011; Acevedo Garcia et al. 2014; Kusch et al. 2016; Polanco et al. 2018). However, clade IV harbors MLOs involved in PM susceptibility in monocots. For example, *HvMLO* in barley and *TaMLO1-A1*, *TaMLO1-B1*, and *TaMLO1-D1* in wheat are found in clade IV (Wang et al. 2014; Acevedo Garcia et al. 2014). Therefore, TaMLOs found in clade IV (*TaMLO3/6-A1, TaMLO3/6-A2, TaMLO3/6-A3, TaMLO3/6-B1, TaMLO3/6-B2, TaMLO3/6-D1,* and *TaMLO3/6-D2*) are possibly associated with PM susceptibility. These genes are the potential targets for antisense expression and the CRISPR/Cas9 system to confer PM resistance in wheat.

### 3.4. Gene Structure and Conserved Domain/Motif Analysis

#### 3.4.1. Comparison of Gene Structure and Motifs with Phylogenetic Tree

The exons and introns are of great importance for plant biodiversity, variation among gene family members, and gene functions. The gene structure analysis revealed that TaMLO genes have 9 to 15 exons. A total of 21 (45%) harbor 13 exons, while 12 genes have 14 exons. Similarly, 15 and 11 exons are present in 5 genes each. Only two genes have 10 exons (*TaMLO1-A1* and *TaMLO12-A1*), and 9 exons are present in *TaMLO12-B1* and *TaMLO12-D1*. It was also noticed that 6 genes have no upstream and downstream regions, 3 genes are without upstream regions, and 2 genes have no downstream regions (**Figure 2b**). This range of exon distribution for MLO genes is well conserved in many plant species. For example, there were 8 to 17 exons in bitter gourd (Chen et al. 2021), 12 to 15 exons in *B. distachyon* (Ablazov and Tombuloglu 2016), 12 to 16 exons in several Rosaceae species (Tian et al. 2022), 13 to 15 exons in lentil (Polanco et al. 2018), and 12 to 17 exons in chickpea, mung bean, pea, groundnut, narrow-leaf lupin, pigeon pea, and common bean (Rispail and Rubiales 2016). All members of groups I, III, and IV in the phylogenetic tree have distinct intron/exon distributions, while group II shows three different intron/exon patterns, which are consistent with the division of group II into three subgroups. Similarly, motif distribution in group II also has three distinct patterns, which should be responsible for its classification into three subgroups. Taken together, the diversity in gene structure and motifs is strongly associated with clade formation in phylogenetic trees.

#### 3.4.2. Characteristics of Conserved Domains/Motifs in MLO Genes

The MEME server identified 16 conserved motifs in 47 MLO genes with a width range of 11 (motif 12) to 50 (motif 5) amino acids; the motif sequences are given in **Table S3.** The nature, number, and patterns of motifs in different subfamilies/groups of the phylogenetic tree are diverse but conserved; e.g., there are ten motifs in group I, 11 to 14 motifs in subgroups of group II, 15 motifs in group III, and 12 motifs in group IV. Eight of these 16 motifs were highly conserved; e.g., motifs 1 to 4 are present in all 47 MLO genes, motifs 6 and 7 are present in 46 genes, while motifs 5 and 8 are present in 45 and 42 genes, respectively. It demonstrates the basic structural and functional resemblances among all genes. For further validation, we searched all 16 motifs in the SMART database, which showed that motifs 1 to 8 were an integral part of the MLO domain. And motifs 1, 2, 7, and 8 were transmembrane domains/regions (TMD) (**Table S3; Figure 2c**).

The presence of motifs 1 and 2 in all MLOs and motifs 7 and 8 in 97.87% and 89.36% of MLO proteins indicated that these are hydrophobic amino acids, which anchor the MLO proteins in the cell membrane. MLO proteins were searched on the Deep TMHMM server, which revealed that 46 of 47 MLOs (97.87%) had a seven-transmembrane (TM) structure (**Figure 3; Table S2**), which was a feature of the MLO genes (Devoto et al. 2003; Chen et al. 2009). While only *TaMLO7-A1* had six transmembrane helices, it was still classified as MLO protein due to the presence of the MLO domain. MLOs are a group of plant disease-susceptible genes that are leucine-rich (9.9 to 13.1%) (Ablazov and Tombuloglu 2016). We found that all the 47 wheat MLO genes are leucine-rich (8.5 to 12.9%) as shown in **table S2**, thus predicting their role in pathogen response.

### 3.5. Comparative Phylogeny of wheat, *A. thaliana* and rice MLOs

A comparative phylogenetic tree was constructed by using the protein sequences of 47 TaMLOs, 15 MLOs of *A. thaliana*, and 12 MLOs of rice, which classified the 74 MLOs of three species into four clades (**Figure 4**). The clade numbers were assigned by following the clade numbering in *A. thaliana* (Devoto et al. 2003), as per standard practice for MLOs. The clades I, II, III, and IV harbored 6, 28, 6, and 7 TaMLO proteins, respectively, just like the phylogenetic tree of 47 TaMLO genes (**Figure 2a**). Additionally, the comparative tree also had three sub-clades for clade II, thus showing a conserved pattern of clustering for both the phylogenetic trees (**Figure 4**; **Figure 2a**). Clade I harbored two rice, three Arabidopsis, and six wheat MLOs. The clade II harbored 28 of 47 (59.54%) TaMLOs, 6 of 12 OsMLOs, and three AtMLOs, i.e., *AtMLO1*, *AtMLO13*, and *AtMLO15*. The subclade IIA consisted exclusively of monocots, i.e., Os*MLO5*, *OsMLO10*, and 14 TaMLOs. Clade III had two rice, five Arabidopsis, and six wheat MLOs, while clade IV contained two rice, four Arabidopsis, and seven wheat MLOs (**Figure 4**). Similarly, a comparative tree consisting of rice, barley, maize, bread wheat, *A. thaliana*, *Physcomitrella patens*, tomato, mustard, and pepper MLOs also classified the genes into four clades (Liu and Zhu 2008).

**Figure 4:**
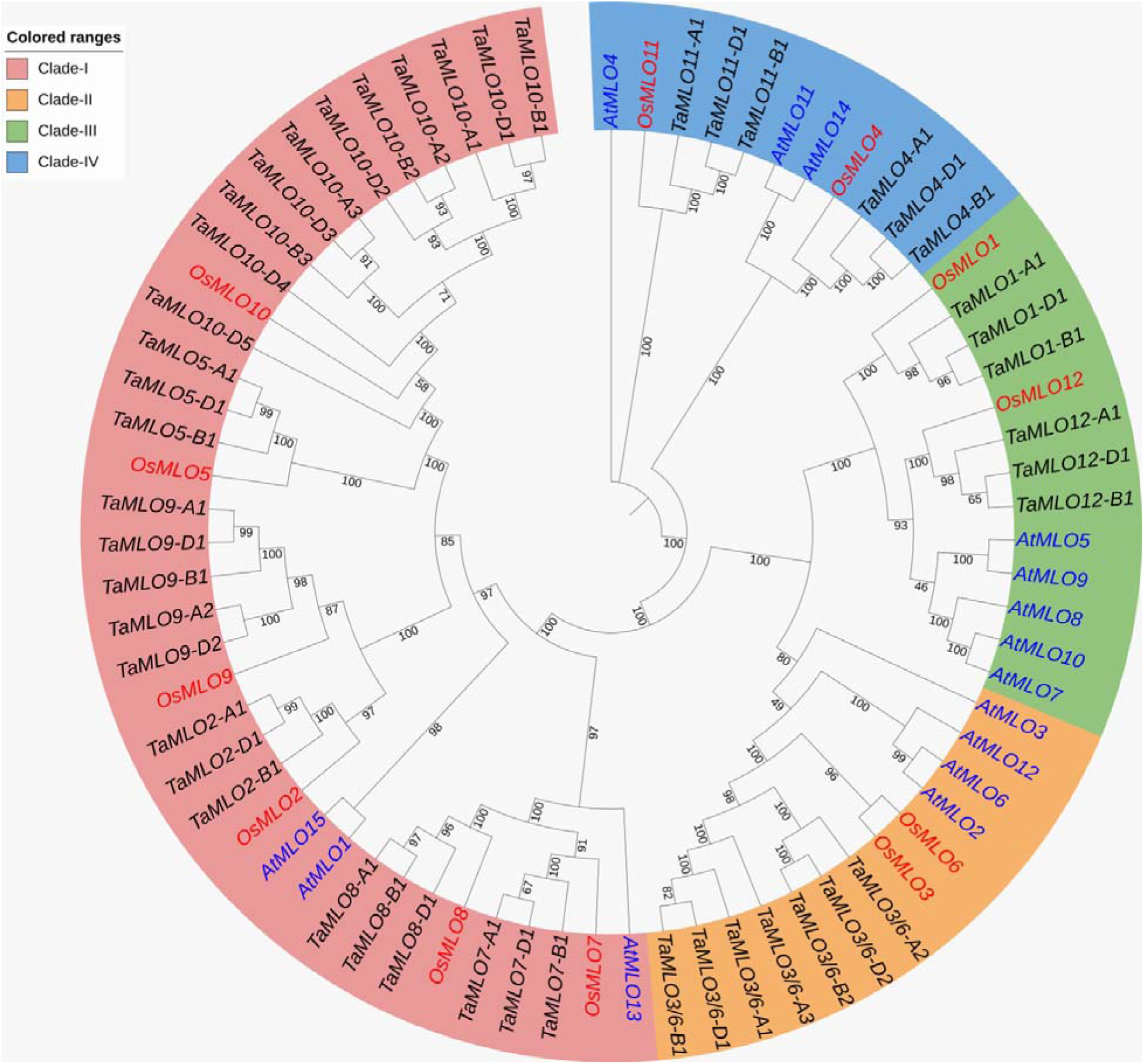
A multi-species comparative phylogenetic analysis of 47, 15, and 12 MLO proteins belonging to bread wheat, *A. thaliana,* and rice, respectively. The tree was constructed by the maximum likelihood method with 1000 bootstrap replications. The numbers at the nodes show the confidence for the branching point, and a high number indicates that the taxa branching from a specific node are closely related. The classification pattern showed four clades, and wheat, *A. thaliana*, and rice genes are represented by black, red, and blue colors, respectively.

However, most of the comparative phylogenetic trees do not support this pattern. Several comparative trees classified the MLOs from rice, wheat, *B. distachyon*, barley, maize, *A. thaliana*, pea, tomato, pepper, wax gourd, bottle gourd, melon, cucumber, watermelon, pumpkin, crookneck pumpkin, kaffir lime, peach, apricot, Chinese white pear, wild strawberry, black raspberry, and apple into six distinct clades (Konishi et al. 2010; Ablazov and Tombuloglu 2016; Tian et al. 2022; Gong et al. 2025). Likewise, some comparative trees based on legume (chickpea, lentil, barrel medic, pea, common bean, narrow-leaved lupin, pigeon pea, groundnut, soybean, and mung bean) MLOs classified the genes into seven distinct clades (Rispail and Rubiales 2016; Polanco et al. 2018), while clade VII is specific to legumes. Additionally, comparative trees based on wheat, maize, barley, rice, *A. thaliana*, pepper, cucumber, tomato, pea, apple, peach, tobacco, and barrel clover MLOs showed eight distinct clades (Zheng et al. 2016; Tapia et al. 2021). This consistent occurrence of six to eight clades across diverse plant species suggests a deep-rooted and robust evolutionary framework for MLO genes.

The distribution of MLOs into different clades is associated with specific functions of these genes. For example, clade I members, *AtMLO4* and *AtMLO11*, are involved in growth behavior or root thigmomorphogenesis (root curling responding to tactile stimulus) (Chen et al. 2009). Additionally, a clade III gene, *AtMLO7*, is involved in pollen tube perception of the egg cell (Kessler et al. 2010), and *OsMLO12* is involved in pollen hydration or reproduction (Yi et al. 2014). Furthermore, clade IV members, i.e., *AtMLO2*, *AtMLO6*, *AtMLO12,* barley *HvMLO*, and three wheat genes, i.e., *TaMLO-A1*, *TaMLO-B1*, and *TaMLO-D1* (renamed as *TaMLO3/6-A2*, *TaMLO3/6-B2*, and TaMLO3/6-D2), are involved in PM susceptibility (Acevedo Garcia et al. 2014). However, *AtMLO2*, *AtMLO6*, and *AtMLO12* are classified as clade V members for *A. thaliana*, as clade IV was lost in Arabidopsis during the evolution (Kusch et al. 2016).

However, this classification is being debated and challenged, and several genes have quite different functions as compared to their location in clades. For example, a clade I member, *AtMLO14*, is not involved in root thigmomorphogenesis, unlike other clade I members, i.e., *AtMLO4* and *AtMLO11* (Chen et al. 2009). Similarly, members of clade III, such as peach *PpMLO1* and *AtMLO7*, are also involved in susceptibility to PM invasion (Kessler et al. 2010; Acevedo Garcia et al. 2014). Additionally, a clade II member, *OsMLO2*, was also involved in PM susceptibility (Acevedo-Garcia et al. 2017). This situation becomes further complicated by the fact that MLOs are classified into six, seven, or eight clades in different species.

Therefore, it can be concluded that in addition to clade IV MLOs (*TaMLO3/6-A1*, *TaMLO3/6-A2*, *TaMLO3/6-A3*, *TaMLO3/6-B1*, *TaMLO3/6-B2*, *TaMLO3/6-D1*, *TaMLO3/6-D2*), some clade III members (*TaMLO12-A1*, *TaMLO12-B1*, *TaMLO12-D1*), i.e., orthologs of *AtMLO7*, could also be involved in PM susceptibility. Similarly, wheat orthologs (*TaMLO2-A1*, *TaMLO2-B1*, *TaMLO2-D1*) of the rice PM susceptibility gene, *OsMLO2*, could also be involved in PM susceptibility, despite being part of clade II. Interestingly, not even a single gene clustered with *AtMLO3*, as this gene has been classified as a member of clade VI (Kusch et al. 2016). In conclusion, comparative phylogenetic analysis provided valuable insights into the evolutionary patterns and functional implications of these MLO proteins.

### 3.6. Gene Duplication and Evolution Analysis of TaMLOs

The CDS of 47 TaMLO genes were MAFFT aligned (pairwise) in all possible combinations by Sequence Demarcation Tool v1.2 (Muhire et al. 2014). The alignments with ≥90% sequence homologies were considered to be duplicated genes. Gene duplication is an essential pathway for the evolution of plant gene families. A total of 50 duplicated gene pairs were present in wheat, which involved 42 TaMLO genes. A total of 94% (47) of duplications were segmental, i.e., duplicated genes occurred on different chromosomes (**Table S4**). Thus, segmental duplications are prevalent in wheat MLOs. It was also reported for the MADS-Box gene family in wheat (Raza et al. 2022) and the APETALA-2 gene family in rice (Ahmed et al. 2021). The Thus, segmental duplications are more prevalent in monocots, e.g., 94% in wheat. On the other hand, dicot tomato has 36.2% segmental duplications (Shi et al. 2020). The remaining 6% duplications were tandem. The gene pairs with tandem duplications formed gene clusters on chromosomes 2A, 2B, and 2D (**Table S4**).

Further evolution analysis was performed by calculating the synonymous substitution rate (Ks) and non-synonymous substitution rate (Ka) per site per year for all duplicated MLO gene pairs. We used the values of Ks to calculate the time of divergence/duplication for TaMLO duplicated pairs, and it was found to be between 3.60 and 19.83 million years ago (MYA) (**Table S4**). The Ka/Ks ratios measure the selection pressure on amino-acid substitutions during the evolution. The synonymous mutations are not known to alter the amino acid sequences, but most of the non-synonymous mutations are deleterious. Therefore, the base substitution rates are very low under purifying selection. The values of Ka/Ks ranged between 0.052 and 0.373; as Ka/Ks < 1, it implies that synonymous substitutions are more prevalent as compared to non-synonymous ones (**Table S4**). The Ka/Ks < 1 indicates that duplicated genes have negative or purifying selection. The purifying selection was also reported for MLOs in *B. distachyon*, rice, maize, *A. thaliana*, cucumber, tomato, melon, grapevine, Amborella, several Rosaceae species, and black cottonwood (Liu and Zhu 2008; Shi et al. 2020; Tian et al. 2022; Gong et al. 2025). It can be concluded that many plants have employed purifying selection to remove deleterious alleles during evolution.

### 3.7. MicroRNAs (miRNAs) Potentially Targeting the Wheat MLOs

The miRNAs targeting the wheat MLOs were identified through the psRNATarget server by using the transcripts of MLOs. We identified a total of 121 miRNAs targeting 41 wheat MLO genes, including eight miRNAs targeting the *TaMLO3/6-B2* gene. Likewise, seven miRNAs each targeted the *TaMLO3/6-A1* and *TaMLO4-D1* genes, while six miRNAs targeted the *TaMLO1-B1* gene. Ttae-miR5384-3p, tae-miR9679-5p, tae-miR9657a-3p, tae-miR9781, tae-miR167c-5p, and tae-miR167a are the most prevalent miRNAs that target the MLOs. The miRNAs, their size and sequences, and target regions are given in **table S5**. The miRNAs are 19-24 nt small RNA molecules that bind with the mRNA of a gene by complementary pairing and either cleave the target or repress its translation, thus reducing its expression (Budak et al. 2015). Several miRNAs are involved in natural stress response in plants by targeting the stress-related genes at transcription and post-transcription levels. Therefore, miRNAs identified in this study can be overexpressed in wheat to silence the MLO genes to confer PM resistance.

3.8. *In silico* Expression Analysis of MLO Genes

#### 3.8.1. MLOs Expression at Different Stages of Development

MLO gene expression patterns have been extensively investigated in crop plants during growth and development. In this study, we analyzed TaMLO expression perturbations in several tissues at different time points to get insights into functional roles under prevalent conditions. Out of a total 47 genes, 28 (∼60%) were expressed in at least two different tissues during the whole growth and development period (**Figure 5**). Among these, at least 9 genes (*TaMLO9-A1, TaMLO10-A1, TaMLO3/6-A2, TaMLO8-A1, TaMLO3/6-B2, TaMLO8-B1, TaMLO3/6-D2, TaMLO8-D1*, and *TaMLO10-D4*) showed ubiquitous expression in nearly all investigated tissues. Interestingly, all these genes exhibited highly significant expression patterns in the flag leaf sheath and blade during the reproductive phase (30% spike to grain ripening stages). However, several other genes exhibited tissue-specific and growth-stage-specific expression perturbations (**Figure 5**). Collectively, these expression patterns indicate diverse functions of MLO genes in wheat growth and development. The MLO proteins are well documented to be involved in growth and development processes such as pollen tube reception in carpel (Kessler et al. 2010), root thigmomorphogenesis (Chen et al. 2009; Bindzinski et al. 2014), reproductive development (Davis et al. 2017) in Arabidopsis, and pollen hydration in rice (Yi et al. 2014). Similarly, a systematic study of rice MLO genes indicated their roles in morphological development, such as the formation of stigma and root tips, in addition to photoperiod sensitivity (Nguyen et al. 2016). Thus, it can be concluded that MLOs also have a role in plant growth and development in addition to immunity.

**Figure 5:**
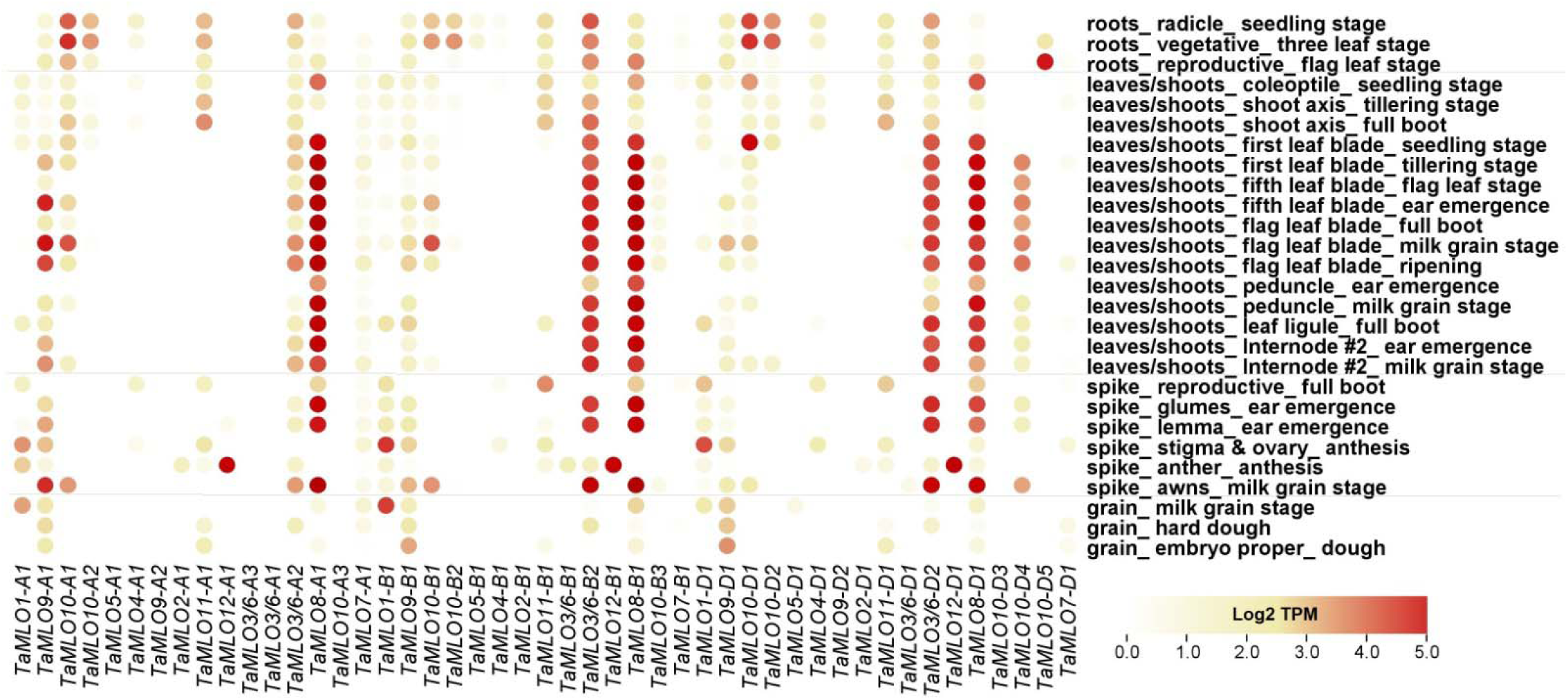
*In silico* expression patterns of 47 TaMLOs during different stages of growth and development in wheat.

#### 3.8.2. MLOs Expression in Different Abiotic Stresses

We also studied the transcriptional changes in wheat MLOs under four different abiotic stresses, including cold, drought, heat, and phosphate starvation. Almost ≥60% of genes were expressed after at least one abiotic and/or biotic stress application (**Figure 6**). Interestingly, all those genes that were highly ubiquitously expressed during different growth and development stages were also highly expressed under abiotic and biotic stresses. Under abiotic stresses, seven genes after two weeks of cold stress (*TaMLO1-A1, TaMLO3/6-A2, TaMLO9-B1, TaMLO1-D1, TaMLO9-D1, TaMLO10-D1*, and *TaMLO10-D4*), six genes after 1–12 hours of drought stress (*TaMLO10-A2, TaMLO10-B1, TaMLO10-B2, TaMLO9-D1, TaMLO10-D1*, and *TaMLO10-D2*), three genes after 1–6 hours of heat stress (*TaMLO10-B3, TaMLO10-D1*, and *TaMLO10-D3*), and three genes under phosphate starvation at the vegetative stage (*TaMLO3/6-A2, TaMLO10-B2*, and *TaMLO10-D1*) exhibited substantial transcriptional perturbations compared with their respective control (CK) treatments (**Figure 5**). Contrary to this, none of the seven wheat MLOs showed differential expression under salt and osmotic stress (Konishi et al. 2010) despite expression under salinity. However, *AtMLO4, 6, 8, 11, 12*, and *13* genes were upregulated, and *AtMLO10, AtMLO15,* Os*MLO10,* and *OsMLO12* were downregulated under salinity (Chen et al. 2006; Nguyen et al. 2016). Similarly, *BdMLO2*, *BdMLO7, OsMLO3*, *OsMLO4*, *OsMLO7*, *OsMLO8*, *OsMLO9*, *OsMLO10*, and *OsMLO12* genes (Ablazov and Tombuloglu 2016; Nguyen et al. 2016) showed differential expression under drought stress. Likewise, *AtMLO8*, *AtMLO12*, *BdMLO2, OsMLO1, OsMLO3, OsMLO4,* and *OsMLO10* genes were upregulated, while *AtMLO2*, *AtMLO7*, *BdMLO6*, 7, 8, 9, and 10 were downregulated (Chen et al. 2006; Ablazov and Tombuloglu 2016; Nguyen et al. 2016) under cold stress. Under heat stress, *OsMLO5*, *7, 9,* and *10* were upregulated, while *OsMLO1* was downregulated (Nguyen et al. 2016). However, the role of MLOs under phosphate starvation has not been studied yet. It can be concluded that there is an emerging role for MLO genes in abiotic stresses that can be used to develop climate-smart plants.

**Figure 6:**
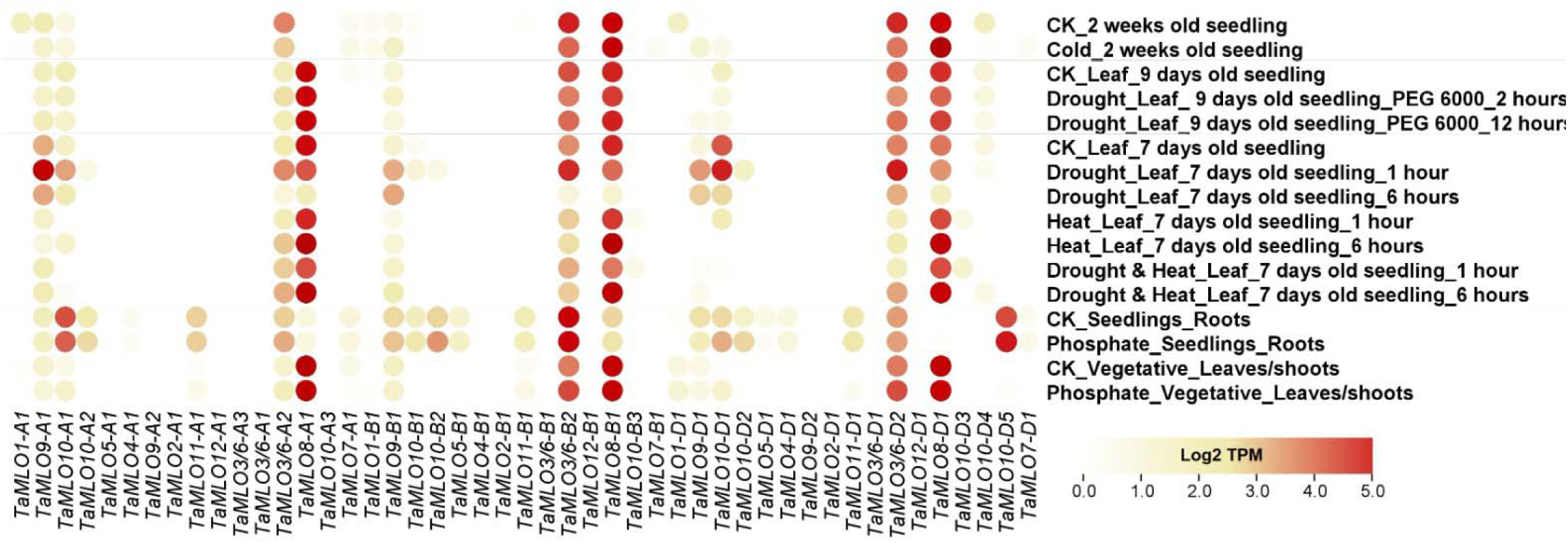
*In silico* expression patterns of 47 TaMLOs under different abiotic stresses in wheat.

#### 3.8.3. *In silico* MLOs Expression under Different Biotic Stresses

We also studied the expression of TaMLOs under three different biotic stresses, including fusarium head blight (FHB), PM, and stripe rust (SR) inoculations. Under biotic stresses, at least four genes (*TaMLO10-A2, TaMLO3/6-A2, TaMLO7-A1,* and *TaMLO10-D5*) exhibited PM-specific transcriptional perturbation when compared with their respective control treatment. Whereas, only *TaMLO9-D1* showed stripe rust-specific expression changes. Likewise, at least five genes (*TaMLO10-A2, TaMLO4-A1, TaMLO10-B1, TaMLO10-B2, TaMLO10-D1*, and *TaMLO10-D2*) exhibited FHB-specific expression patterns when compared with respective control treatments (**Figure 7)**.

**Figure 7:**
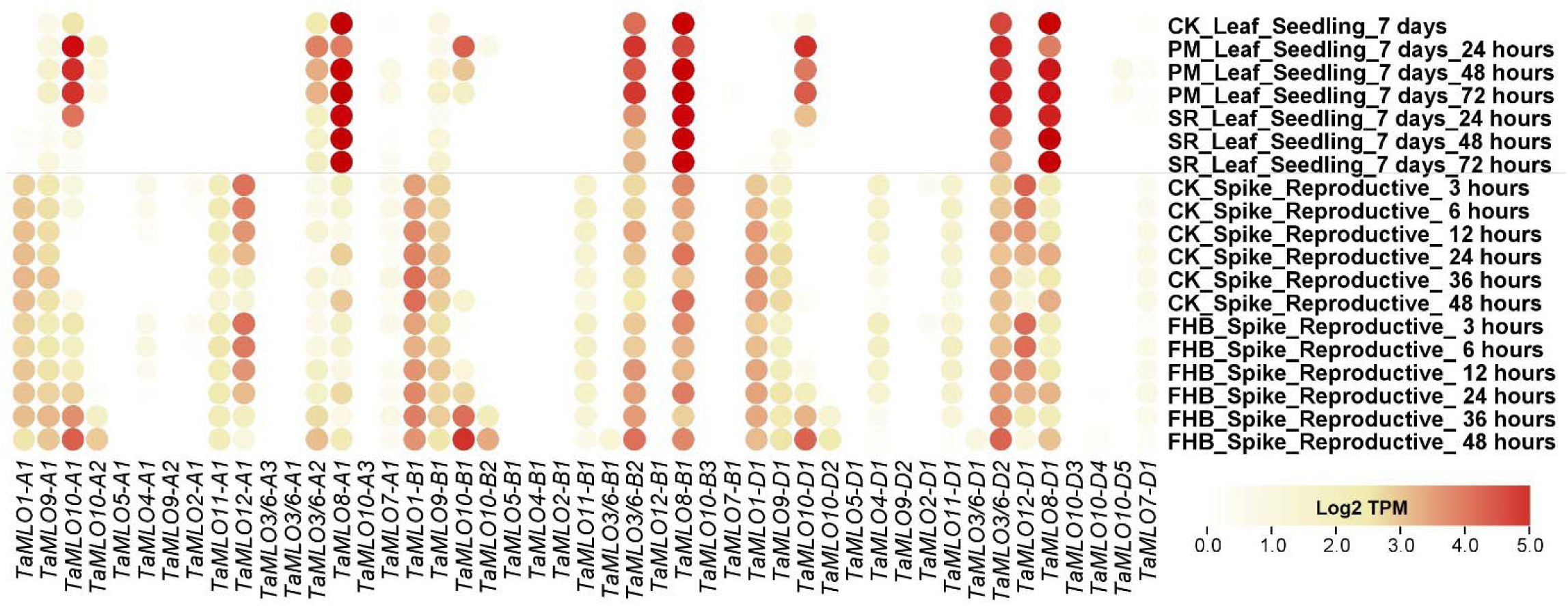
*In silico* expression patterns of 47 TaMLOs under different biotic stresses in wheat.

MLO family genes are unique to plants, governing resistance against PM infections through a negative regulatory pathway. However, apart from their role in wheat PM susceptibility, they have roles in resistance/susceptibility against wheat stripe rust (Zhang et al. 2014), FHB (Ablazov and Tombuloglu 2016; Acevedo-Garcia et al. 2017), rice blast fungus (Nguyen et al. 2016), *Pseudomonas syringae* (Acevedo-Garcia et al. 2017), and several bacterial and oomycete pathogens (Acevedo Garcia et al. 2014). In this study, a substantial number of MLOs also showed ubiquitous expression patterns during growth and development and abiotic and biotic stresses. These results support previous findings that these plant-specific genes regulate multifarious processes for successful completion of the wheat plant life cycle and survival under unfavorable conditions. Detailed functional characterization of significantly perturbed MLO genes using modern molecular and biotechnological techniques such as CRISPR-Cas9 could yield climate-resilient and super-immune wheat cultivars.

### 3.9. Quantitative RT-PCR Analysis of TaMLO Genes

PM is one of the most devastating diseases of wheat grown in temperate regions. Several studies have reported gene expression analyses of wheat genes following *Bgt* inoculation. For example, a large-scale transcriptome analysis of the Bgt resistant line revealed distinct expression patterns responding to PM and stripe rust in wheat (Zhang et al. 2014). Similarly, a gene co-expression network analysis provided novel insights into the dynamic response of wheat to PM resistance (Hu et al. 2020). Since our *in-silico* expression analysis exhibited higher expressions of several genes during growth & development and under abiotic and biotic stresses. We selected 10 genes with overlapping patterns under these conditions for qRT-PCR expression validation under PM infection. In general, expressions of these genes started to elevate 24 hours post infection (hpi) and reached peak deregulations at 72 hpi (**Figure 8**).

**Figure 8:**
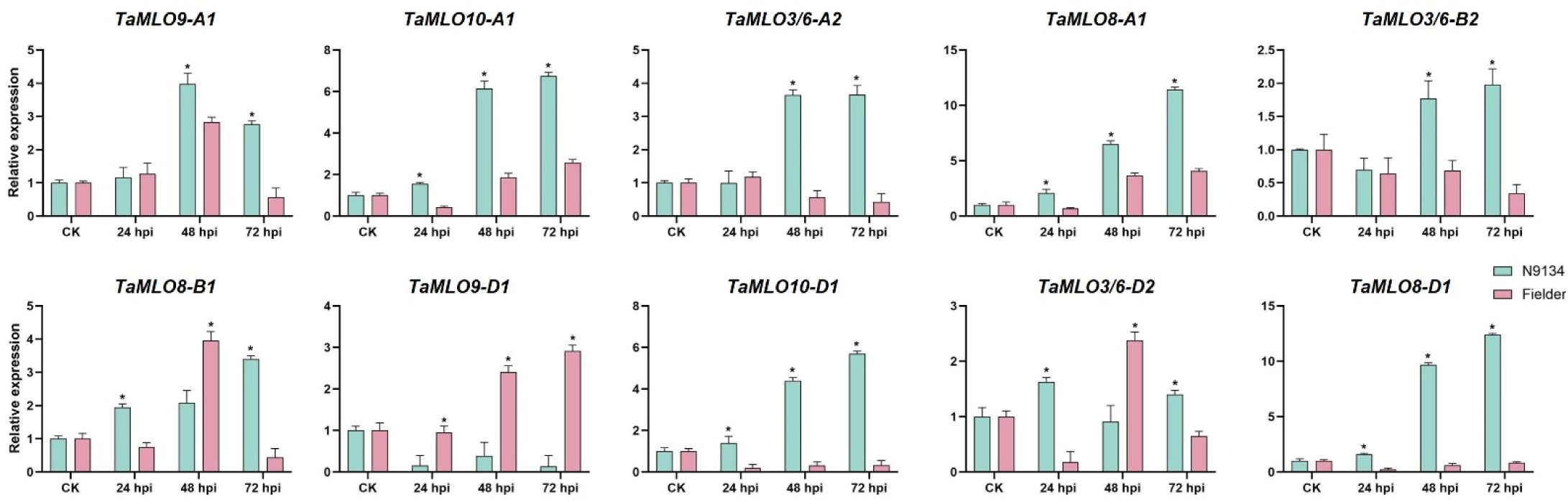
qRT-PCR based expression validation of wheat MLO genes. The relative expressions were quantified after 24, 48 and 72 hours of post inoculation (hpi) with *Bgt*. Student’s t-test was used to compare relative expression (*P* < 0.05) between PM resistant (N9134) and susceptible (Fielder) wheat genotypes.

Homeologs of *TaMLO8*, *TaMLO10,* and *TaMLO3/6* exhibited significant upregulation in the *Bgt*-resistant wheat line (N9134) as compared with the susceptible wheat cultivar (Fielder). However, *TaMLO9-A1* exhibited upregulation, whereas *TaMLO9-D1* showed downregulation in the *Bgt*-resistant line. The qRT-PCR-based expressions were consistent with *in silico* expression patterns. Four genes (*TaMLO8-A1*, *TaMLO8-D1*, *TaMLO10-A1,* and *TaMLO10-D1*) exhibited significant upregulation in the resistant line after 24, 48, and 72 hpi with the Bgt race as compared with the susceptible cultivar. Likewise, five genes (*TaMLO9-A1*, *TaMLO3/6-A2*, *TaMLO3/6-B2*, *TaMLO8-B1*, *TaMLO3/6-D2,* and *TaMLO8-D1*) exhibited significantly higher expression in the resistant line after 2-3 days hpi, indicating positive regulation of PM resistance. Whereas a single gene (*TaMLO9-D1*) showed downregulation in the resistant wheat line, suggesting a negative regulation of PM resistance **(Figure 8)**.

Notably, seven of these genes belonged to clade II, whereas 3 genes belonged to clade IV, and all shared homeologous relationships among distinct clades. These genes are expected to be involved in PM susceptibility, and are candidate to be knocked out to confer PM resistance. Indeed, three of these clade IV genes (*TaMLO3/6-A2*, *TaMLO3/6-B2*, and *TaMLO3/6-D2*) have been knockout out to confer durable and stable PM resistance in wheat. For example, simultaneous knock out of *TaMLO-A1* (renamed as *TaMLO3/6-A2)*, *TaMLO-B1* (renamed as *TaMLO3/6-B2),* and *TaMLO-D1* (renamed as *TaMLO3/6-D2)* genes by TALEN, TILLING, and CRISPR/Cas9 in bread wheat conferred heritable broad-spectrum PM resistance without pleiotropic phenotypes (Wang et al. 2014; Acevedo Garcia et al. 2017). Recently, CRISPR/Cas9-based knockout of *TaMLO-B1* in wheat that resulted in robust PM resistance without any pleiotropic effects (Li et al. 2022). Additionally, simultaneous mutations in these three genes including some weak mutations in one of these genes provided highly effective PM resistance without pleiotropic effects (Ingvardsen et al. 2023). Interestingly, none of the clade II genes have been characterized in any plant species yet. However, as the above mentioned seven MLOs showed differential expression under PM resistance, their involvement in PM susceptibility is very probable. As we discussed in comparative phylogenetic section, clade I and III genes are also involved in PM susceptibility, so we believe that MLO genes belonging to all four clades in monocots are involved in PM susceptibility.

Notably, wheat MLO genes have been reported to confer resistance against stripe rust and wheat blast (Zhang et al. 2014; O’Hara et al. 2024). In this study, we also observed overlapping expression patterns of several genes under different abiotic and biotic stresses, as well as during critical growth and development stages. These data indicate involvement of identified wheat MLO genes in conferring broad-spectrum resistance against different stresses. In future, it would be interesting to clone and edit these genes for engineering a durable PM resistance in wheat cultivars.

## Author Contributions

**BH**: Experiment design, analysis, writing original draft. **QR**: Analysis, experimentation, writing original draft. **HR**: Analysis, writing original draft. **RMA**: Conceptualization, experiment design, supervision. **MFA**: Analysis, editing of draft. **MY**: Editing of draft. **HB**: Supervision, editing of draft. **ZA**: Experiment design, supervision, provided resources for qRT-PCR. All authors reviewed the manuscript.

## Availability of Data and Materials

No datasets were generated during the current study.

## Declarations

### Ethics Approval and Consent to Participate

Not applicable.

### Consent for Publication

Not applicable.

